# Metabolome-informed microbiome analysis refines metadata classifications and reveals unexpected medication transfer in captive cheetahs

**DOI:** 10.1101/790063

**Authors:** Julia M. Gauglitz, James T. Morton, Anupriya Tripathi, Shalisa Hansen, Michele Gaffney, Carolina Carpenter, Kelly C. Weldon, Riya Shah, Amy Parampil, Andrea Fidgett, Austin D. Swafford, Rob Knight, Pieter C. Dorrestein

## Abstract

Even high-quality collection and reporting of study metadata in microbiome studies can lead to various forms of inadvertently missing or mischaracterized information that can alter the interpretation or outcome of the studies, especially with non-model organisms. Metabolomic profiling of fecal microbiome samples can provide empirical insight into unanticipated confounding factors that are not possible to obtain even from detailed care records. We illustrate this point using data from cheetahs from the San Diego Zoo Safari Park. The metabolomic characterization indicated that one cheetah had to be moved from the non-antibiotic-exposed to the antibiotic-exposed group. The detection of the antibiotic in this second cheetah was likely due to grooming interactions with the cheetah that was administered antibiotics. Similarly, because transit time for stool is variable, early fecal samples within the first few days of antibiotic prescription do not all contain detectable antibiotics. Therefore, the microbiome is not affected by the antibiotics at those time points. These insights significantly altered the way the samples were grouped for analysis (antibiotic vs no antibiotic), and the subsequent understanding of the effect of the antibiotics on the cheetah microbiome. Metabolomics also revealed information about numerous other medications, and provided unexpected dietary insights that in turn improved our understanding of the molecular patterns on the impact on the community microbial structure. These results suggest that untargeted metabolomics data provide empirical evidence to correct records of non-model organisms in captivity, although we also expect these methods will be appropriate for experimental conditions typical in human studies.

**Importance:** Metabolome-informed analyses can enhance ‘omics studies by enabling the correct partitioning of samples by identifying hidden confounders inadvertently misrepresented or omitted from carefully curated metadata. We demonstrate the utility of metabolomics in a study characterizing the microbiome associated with liver disease in cheetahs. Metabolome-informed reinterpretation of metagenome and metabolome profiles factored in an unexpected transfer of antibiotics preventing misinterpretation of the data. Our work suggests that untargeted metabolomics can be used to verify, augment, and correct sample metadata to support improved grouping of sample data for microbiome analyses, here for non-model organisms in captivity. However, the techniques also suggest a path forward for correcting clinical information in human studies to enable higher-precision analyses.

## 1. Introduction

The microbiome is accepted as a critical aspect of organismal health, with much attention being focused on the gut microbiome. This environment is variable, in part defined by an irregular flow of inputs, including diet and medications such as antibiotics, that impact and shape the microbial community (1–7). Common ways to examine this variability are to profile microbial community structure and functional capacity, and less frequently functional activity and output, including metabolite signatures. Correct interpretation of these profiles relies on the detailed, relevant metadata that form the foundation for all analyses and applications of the data.

A mismatch between collected metadata and variables of interest can result from a multitude of reasons, with reporting errors and biases, omission of categories in the metadata based on original study design, unexpected exposures, and poorly structured or inconsistent descriptors amongst the most common. The approach of some large human cohorts, such as the American Gut Project (8), has been to capture an extensive array of information using controlled vocabulary and values to maximize the likelihood of having relevant metadata. However, cohorts that rely on self-reported information and self-initiative to complete run the risk of obtaining erroneous, incomplete, and variable amounts of metadata for each sample. Social pressures or fear of repercussions may further prevent the disclosure of illicit or sensitive information such as drug use, sexually transmitted diseases, poor hygiene or diet, etc. and temporal distance inevitably complicates the recall of dietary, medication, and health-related events (9). For example, metabolite analysis detected the presence of antibiotics in fecal samples from individuals that reported not having taken antibiotics in the past 6 months or more (8). Furthermore, a central issue across microbiome studies is that it is very challenging to capture additional participant information after a study has been completed, either due to the self-reported nature, unresponsive subjects, or simply the passage of time.

We propose that metabolite-informed microbiome analyses, where the small-molecule composition of a sample, readily detected using a liquid chromatography - tandem mass spectrometry (LC-MS/MS) workflow, can be used to generate empirically determined metadata assignments for compounds such as medications, including antibiotics or painkillers, and personal care products, such as sunscreen. In particular, the use of antibiotics in a clinical setting is common, impacting organisms from livestock, domestic animals, captive wildlife, to humans of every age. Antibiotics have a strong documented impact on the gut microbial community (1–5), often resulting in large decreases in alpha diversity. These changes in alpha diversity have the potential to alter microbial community structure and the gut metabolome. Antibiotics have also been shown to influence the gut microbial communities of members of a household where one individual is taking antibiotics (10). The route of impact was, however, not clear, leaving open the possibility of a shift in the microbiome that is microbially or chemically mediated, possibly through transfer of the drug itself between individuals.

Animals in managed care, such as cheetahs (*Acinonyx jubatus*), have many similarities to human patients in a clinical setting. Interventions are only attempted when deemed medically necessary, and, based on individual health history, each animal has a unique combination of housing location, diet, medication use, and environmental exposure. Cheetahs in captivity suffer from higher rates of veno-occlusive disease and gastrointestinal distress than their wild counterparts (11), leading to multiple treatment interventions, such as changes in diet and medications. However, unlike human subjects, zoo animals do not self-report information, and detailed records of food consumption, medication use, health parameters, housing, and behavior are recorded by keepers and trainers to capture these interventions. These detailed metadata and controlled conditions provide an ideal setting for examining the complementarity of a metabolome-informed approach to microbiome analyses.

Here, we present a workflow (Figure 1) for generating study-specific metabolome-informed metadata categories, using cheetahs as a case study, and highlight the value of the approach for generating empirical metadata and reinterpreting the data. Finally, we discuss broader applications of using empirical evidence to correct sample categorization and records and provide concrete examples where this technique is anticipated to have the greatest impact, suggesting areas where metabolomic data should be routinely collected.

**Figure 1.**
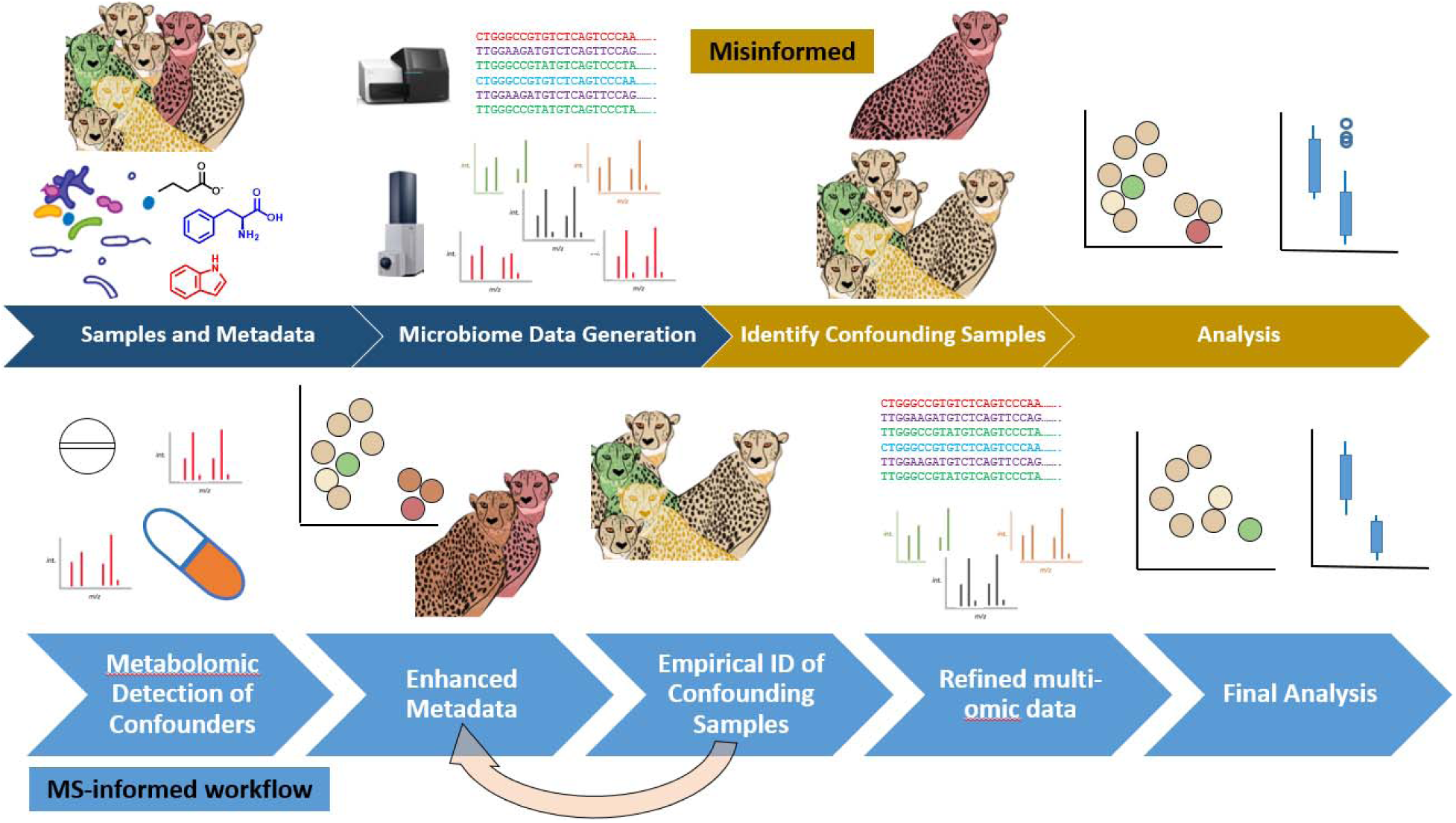
Analysis workflow for metabolome - informed microbiome analyses. All analyses begin with the collection of samples and metadata (information about the samples such as time of collection, participant name or number, etc.), followed by microbiome data generation, such as sequence data and untargeted metabolomics [dark blue]. A traditional, or misinformed, analysis will then identify confounders from the metadata and individually parse the datasets based on reported metadata variables and synthesize results from the datasets in the last analysis step. In contrast, a metabolome-informed analysis, as presented in this study, will empirically determine if there are confounders or additional information identified by metabolomics analysis [see Figure 2], which in this study identified antibiotics in samples where none were reported. The MS data empirically support creation of an MS-informed metadata column, which is applied to each dataset collected (metagenome and metabolome in this analysis) [Figure 3]. This cycle can be iterated over multiple times based on potential confounders and information obtained through the MS analysis. In this case study, a further iteration filtered the metabolite feature table itself, as medication derived metabolites, given only to ill animals, dominated the differences observed [Figure 4]. MS-informed metadata allow the data themselves to help guide the analysis and facilitates communication between the datasets.

## 2. Results

We collected and analyzed paired shallow shotgun metagenomic sequence data and untargeted LC-MS/MS data from fecal samples from seven cheetahs housed at the San Diego Zoo Safari Park in 2018 over the course of 1 month (Table 1). Shallow shotgun metagenomic sequencing (12) provides a snapshot of the microbial community structure at each timepoint and can provide insight into functional potential (e.g., the potential exchange and transformation of metabolites between the environment, microbes, and host). Untargeted LC-MS/MS metabolomics analysis aims to detect all molecules in a sample without any knowledge of the molecular constituents *a priori*, thereby assessing the chemistry of individuals as well as differences between individuals or populations. Broad untargeted metabolomics is able to identify compounds that corroborate or challenge metadata assignments and can thus be inspected in a concerted fashion for accuracy based on individual study design (Figure 1) and therefore provide valuable empirical evidence to enhance the accuracy of our analyses. Combined, these data provide a window into the community structure as well as the inputs that are entering the gut microbial environment, including host, diet, and medication-related compounds.

**Table 1.** Overview of cheetah cohort and selected variables from the metadata. The analyses focus on 7 cheetahs at Wildlife Discoveries. Shotgun metagenomics and metabolomics data are available under Qiita ID 11872 and GNPS ID MSV000082969. Cheetahs maintained the same diet during the 30 day sampling time course. {source: ‘Exp_design_figure.xlsx’ tab2} https://docs.google.com/spreadsheets/d/14mH8m4FgHkGG4J_lTZciJa9kgUNQqWWqGFPQESBM4AY/edit#gid=634406102

| Animal ID | Accession # | Sex | DOB | Age | Liver health | Antibiotic use |
| --- | --- | --- | --- | --- | --- | --- |
| AMARA | 609025 | F | 18-Feb-09 | 9 y 5 mo | Mild disease | no |
| BAHATI | 614426 | F | 1-Sep-14 | 3 y 11 mo | Normal | no |
| JOHARI | 609201 | F | 24-May-09 | 9 y 1 mo | Severe disease | no |
| ISOKA | 615391 | M | 1-Sep-14 | 3 y 11 mo | Normal | yes |
| KIBURI | 610376 | M | 15-Nov-10 | 7 y 8 mo | Normal | no |
| OKUBI | 615390 | M | 1-Sep-14 | 3 y 11 mo | Normal | no |
| RUUXA | 614198 | M | 3-May-14 | 4 y 2 mo | Normal | no |

### 3.1 Discovery of inconsistency between reported and detected MS data

Initial inspection of the metabolome and metagenome identified antibiotic use as a main driver of differences between samples, as observed by principal coordinate analysis (PCoA) (Figure 2a-b), with the first principal axis (PC1) explaining 32.37% of the variance in the metabolome and 70.23% for the microbiome. The between sample distances from PCoA clearly differentiated the majority of the fecal samples belonging to the male cheetah Isoka, who was treated for gastritis with medication, including two 14-day courses of the antibiotic amoxicillin. Interestingly, a fraction of samples reported in the metadata as ‘no antibiotic use’ cluster together with the ‘antibiotic use’ samples, indicating a molecular as well as microbial similarity.

**Figure 2.**
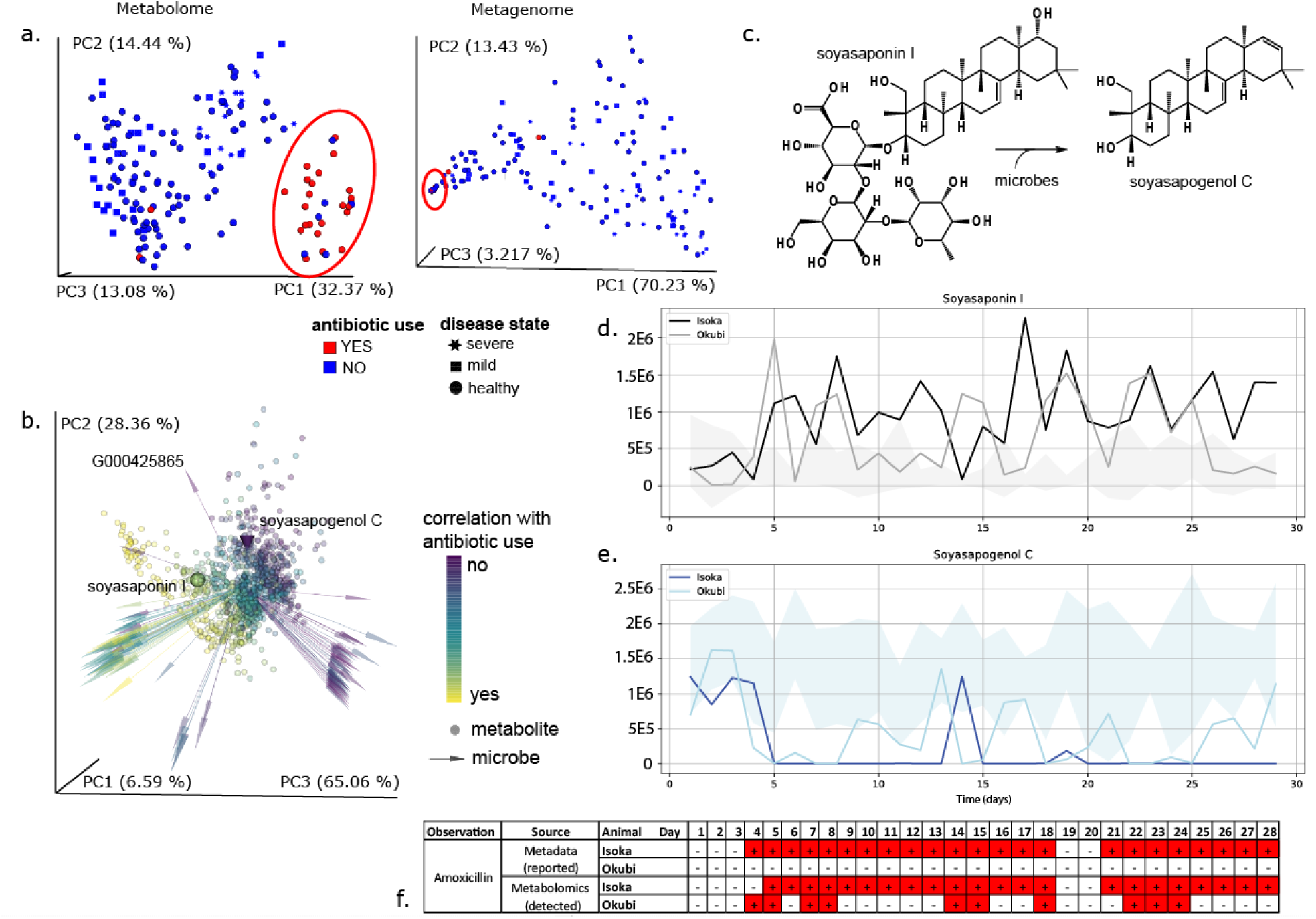
MIM_Fig2_new.ai Antibiotics are a major driver of metabolome and metagenome beta diversity and metabolome analysis reveals unexpected antibiotic transfer, corroborated by differential microbially mediated food metabolism. **a)** Principal coordinate analysis (weighted Unifrac) for MS feature abundance (left) and shotgun sequence data (right) for fecal samples from 7 cheetahs housed at Wildlife Discoveries. Data points are colored by reported antibiotic use based on initial metadata (blue: no; red: yes). Shape designates the disease state, with regards to cheetah liver necrosis syndrome (CLNS). Left is metabolome, right is metagenome; distance metric is weighted Unifrac. **b.** Microbe-metabolite co-occurrence analysis. Large cone represents soyasapogenol C, large sphere represents soyasaponin I. Biplot of cheetah data points - spheres are metabolites, arrows are microbes. Both metabolites and microbes are colored by the same scale based on differential abundance analysis: yellow is associated with antibiotics; purple is not associated with antibiotics. Top 100 species from differential abundance analysis are displayed in the plot; all metabolites are shown. Web of life genome ID: G000425865; NCBI taxonomy: k__Bacteria; p__Firmicutes; c_Bacilli; o_Lactobacillales; f__Carnobacteriaceae; g__*Lacticigenium*; s__*Lacticigenium naphtae*; **c.** Microbially mediated conversion of soyasaponin I to soyasapogenol C; **d.** Soyasaponin I abundance over time for Isoka (black), Okubi (gray) and the range of values for the 5 cheetahs (Amara, Bahati, Johari, Kiburi, and Makena) with no detected amoxicillin (shaded gray); **e.** Soyasapogenol C abundance over time, as plotted in d. Soyasapogenol C is consistently more abundant in feces than soyasaponin I; **f.** Reported amoxicillin administration for Isoka by sampling day (reported), compared with detection of amoxicillin in fecal samples from Isoka and Okubi (detected). Antibiotic prescription or detection are highlighted in red. Days line up with the plots of feature abundance for soyasaponin I and soyasapogenol C. Feature data: soyasaponin I (m/z 441.3731, RT 7.3547 min, row ID 209, annotation network number n/z, correlation group ID 79); soyasapogenol C (m/z 943.5270, RT 5.3381 min, row ID 578, annotation network number 60 correlation group ID 58).

We traced the origin of these ‘no antibiotic use’ samples to one individual, the male cheetah Okubi (Fig 2a-b, blue within the red oval) who did not receive any medications during the sampling period. However, Isoka and Okubi are co-housed male siblings, which presented a risk for misidentification of the source animal for fecal samples, which would impact the findings directly. Therefore, medications were individually orally administered and glitter was added to Isoka’s diet (Rx antibiotic), in order to verify which stool sample originated from which animal. Unlike captive rodents and some domestic animals, cheetahs are not coprophagic, i.e. do not eat each other’s stool, limiting the potential routes of molecular sharing. Instead, we hypothesize that grooming and other social interactions (13) could have led to direct medication carryover.

We tested this hypothesis by examining whether any of the antibiotics Isoka received were detected in the metabolomics data of the fecal samples from Okubi. The hydrolyzed form of amoxicillin was detected by library identification in samples from both Isoka (Rx antibiotic) and Okubi (no Rx) using the web-based global natural products (GNPS) analysis platform (https://gnps.ucsd.edu) (14) which provides Metabolomics Standards Initiative level II or III identifications (15). Moreover, this antibiotic was not detected in any samples from the other cheetahs. During the month long time course, Isoka was on prescription antibiotics for 24 days (out of 29 total samples) and the amoxicillin derivative was detected in 23 samples from Isoka and intermittently in 10 samples from Okubi, which also group together based on PCoA (squares, Figure 2f).

Using a multi-omic microbe-metabolite co-occurrence analysis we observed a strong trend with antibiotic use across both datasets, with highly correlated microbes and metabolites also co-occurring (Figure 2b). Untargeted metabolomics identified members of the molecular family of plant products, soyasaponins, and their degradation products (such as that in Figure 2c) in stool of cheetahs consuming Nebraska Brand Special Beef Feline Diet. These ingredients, not initially part of the metadata collection, were confirmed by the manufacturer ingredient list of the dietary product (http://www.nebraskabrand.com/docs/beefsheet2019pdf.pdf).

One class of correlated molecules includes the soyasaponins, where we identified both precursor (soyasaponin I) and metabolite (soyasapogenol C). Previously, microbial fermentation has been implicated with the conversion of these compounds to their aglycone metabolites (Hu et al., 2004). These results support our findings, as the microbially-derived metabolite soyasapogenol C is correlated with no antibiotic use, while the precursor is correlated with antibiotic use (Figure 2b). In addition to these observed changes in the metabolome following antibiotic exposure, the presence of soyasapogenol C has a high co-occurrence probability with Firmicutes and Proteobacteria, with G000425865 among the most differential microbes in the analysis (Table S1), indicating that changes in microbiota may be driving differences in metabolic function within the gastrointestinal tract.

### 3.2 Metabolome-informed metadata grouping supported by altered microbial metabolism of diet components

Further longitudinal assessment of soyasaponin I and soyasapogenol C (Figure 2d,e) show differential metabolism of soy carbohydrates and mirror the medication schedule for Isoka (Rx antibiotic, Figure 2f) and this empirical evidence supports restructuring of the metadata from antibiotic reported to detected. Antibiotic annotation based on metadata revealed a correlation between antibiotic detection and microbial food metabolism. We observed the breakdown of soyasaponin across animals (Figure 2d, min max range of other animals shaded gray) and a stark absence of the aglycone, soyasapogenol C, when antibiotics were empirically detected in the feces by MS (Figure 2d,e,f).

Fecal samples from Isoka (Rx antibiotic) have a predominance of soyasaponin I (Figure 2d). A longitudinal representation of the relative abundance of these two features for Isoka (Figure 2d,e; black and dark blue) showed an initial ability to metabolize soyasaponin I to soyasapogenol C, which virtually disappeared upon commencing antibiotic treatment. In agreement with the MS-based detection of antibiotics in multiple fecal samples from Okubi (Figure 2f), we also observed a diminished ability to metabolize soyasaponin, which has a stochastic nature, similar to the intermittent observation of antibiotics (Figure 2d,e; light gray and light blue). The remaining cheetahs have higher relative concentrations of soyasapogenol C (Figure 2e, min and max range shaded blue), indicating active microbial deglycosylation.

### 3.3 Empirically derived metadata categorization avoids misinterpretation

Due to the empirical evidence, a new metadata variable was created to delineate the difference between reported and detected antibiotic use, enabling metabolome-informed analyses.

#### Metabolome-informed metabolome analysis

Antibiotics imparted a strong signature to the differences between samples, both in terms of the medication itself, metabolism of soy carbohydrates, as discussed above, as well as changes in host and host-microbial metabolism. The hydrolyzed amoxicillin detected by MS was positively correlated with antibiotic use and highly ranked (Figure 3a), based on a differential abundance analysis. Furthermore, antibiotic use had a profound influence on bile acid metabolism. Conjugated bile acids correlate with antibiotic use (Figure 3b), whereas primary bile acids are negatively correlated with antibiotic use. The log ratio of conjugated (primary and secondary) bile acids divided by primary bile acids is elevated for Isoka and Okubi (Figure 3d) and the difference between the MS-metadata informed categories can be clearly seen in Figure 3c, left and right, respectively. In both cases the difference between antibiotic use and not is statistically significant (Welch’s t-test; p value: 4.5e-49 (MS_reported) and 1.3e-31 (MS_detected). In particular several outliers in no antibiotic use reported are removed when comparing the antibiotic detected categories.

**Figure 3.**
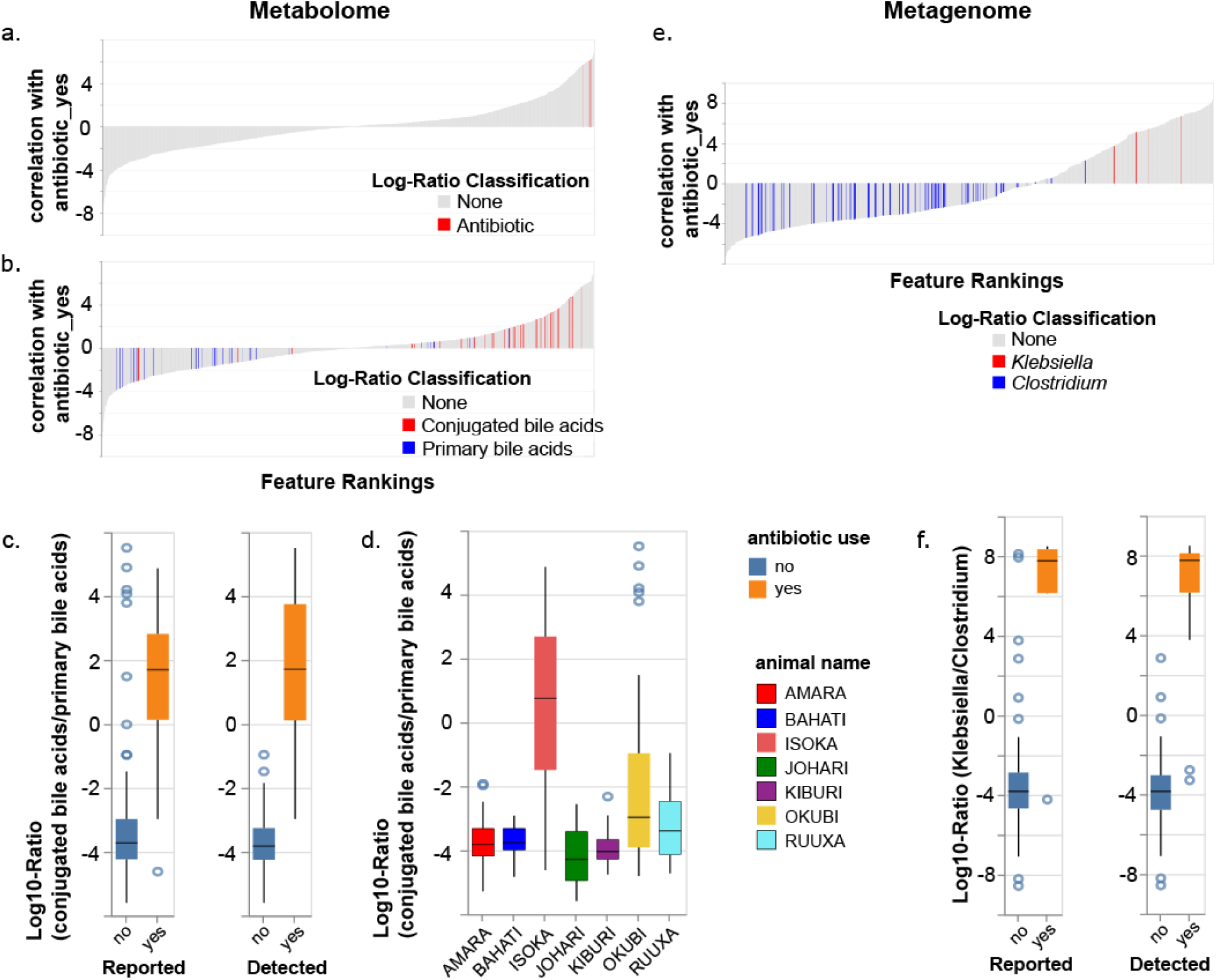
Metabolome data-informed groupings reveal impacts on the metabolome and microbiome profiles. **a-b)** differential abundance analysis of MS features (association with antibiotic yes compared to the antibiotic no as the reference) visualized with Qurro. Positive values in the rank plot correspond to a positive association with antibiotic use. **a.** Antibiotic features highlighted in red; **b.** Impact of antibiotics on bile acid metabolism. Conjugated bile acids in red and primary bile acids in blue. **c-d)** Log ratio of conjugated bile acid features by primary bile acid features, **c.** for reported vs detected antibiotic metadata categorization, respectively, and **d.** plotted by animal. **e.** Differential abundance analysis of MS features (association with antibiotic yes compared to the antibiotic no as the reference) visualized with Qurro. Positive values in the rank plot correspond to a positive association with antibiotic use. Numerator: *Klebsiella* genome IDs (red); Denominator: *Clostridium* genome IDs (blue). **f)** Log ratio of *Klebsiella* to *Clostridium* for reported vs detected antibiotic metadata sorting, respectively. Note the removal of outliers in ‘reported - no’ in ‘detected - no’. The difference between antibiotic use and not for reported and detected in subplots c. and f. are statistically significant (p < 1e-31 for all), based on a Welch’s t-test.

Furthermore, application of this informed metadata variable to the metabolite data still showed that the main driver in separation of cheetah fecal material was antibiotic detection, with PC1 representing 32.37% of the variance in the dataset and effectively separating Isoka and some of Okubi’s samples from the remaining animals (Figure S1a). Based on a PERMANOVA analysis of the weighted unifrac distance matrix, categorization based on metabolite data resulted in a larger effect size (F-statistic for the two different metadata categories: antibiotic use (reported) = 28.2433 compared with antibiotic_presence_WD (detected) = 47.4572.)

Thus, one loop of application of the MS-informed metadata has been completed, applying a new categorization of antibiotic use which reflects the mass spectral data. Recategorization will impact data interpretation: removal of these samples from further analysis reduces the chances of data misinterpretation due to the outliers originally present. Furthermore, the same changes applied based on the MS-informed metadata can be applied to any additional paired dataset.

#### Metabolome-informed metagenome analysis

The metagenome results also have a larger effect size for the metadata assigned due to metabolite analysis (detected) compared with the original metadata (reported) (PERMANOVA F-statistic calculated from the weighted unifrac distance matrix: 41.7599 for antibiotic_use (reported) compared with 75.9906 for antibiotic_presence_WD (detected)) (Figure 1b vs. Figure S1b). The same trend held true for the unweighted distance matrix (Figure S2): F-statistic of 79.4844 for antibiotic_use (reported) compared with 139.961 for antibiotic_presence_WD (detected). Removal of the antibiotic use samples prior to PCoA results in removal of the strong antibiotic signature and reveals a new distribution of samples (Figure S3). Many of Johari’s samples (severe cheetah liver necrosis syndrome (CLNS)) group together in the upper left corner and samples for Amara, with mild CLNS, do not have distinct clustering from the remainder of the healthy fecal samples.

The representation of taxa within metagenomic samples from antibiotic treated animals is likely to be different than healthy controls. We assessed the difference in the taxonomic composition of samples without antibiotic use reported initially (reported) and then with the addition of samples in which antibiotics were detected (detected) (as determined by mass spectral data). There is a stark impact of antibiotic use on the microbial composition of the cheetah fecal samples on antibiotic use (Figure S4, top vs. bottom), which is further differentiated when applying the MS-detected antibiotic grouping (Figure S4, right-hand side). Notably, *Klebsiella* is no longer observable amongst the top 10 most abundant genera when the samples are grouped by antibiotic detection rather than by reporting. *Klebsiella* is also positively associated with antibiotic use, while genera such as *Clostridium* are negatively correlated with antibiotic use (Figure 3e). The log-ratio of *Klebsiella* to *Clostridium* for antibiotic use is higher for antibiotic use in both reported and detected metadata categories (Welch’s t-test; p value 7.7e-61 (detected) and 6.5e-80 (reported)), however notable outliers with a high ratio in antibiotic reported samples in the ‘no’ category are correctly categorized with the application of the metabolome-informed metadata categorization (Figure 3f).

### 3.4 MS-informed refinement of feature table

Cases of metadata inaccuracy due to subjective reporting, missing metadata or unexplained observations have been noted in the literature (8), as well as reported in this manuscript. Regardless of the origin of such observations, it is important to develop strategies to mitigate their impact on the conclusions drawn with regard to the question of interest and empirically derived information allows us to gain insights beyond the information initially collected. MS analysis revealed the presence of S-adenosyl methionine metabolites (from the supplement denamarin) present in the samples from the diseased cheetahs, Amara and Johari, the only ones given the supplement. These features are differentially abundant (Figure 4a), are positively correlated with disease compared with the healthy reference samples. These features, when compared with a log ratio of a general class of all annotated lipids in the metabolite dataset, spread broadly across the rankings (Figure 4a, bottom), are more abundant in samples from Amara and Johari, as compared to the remaining cheetahs, including those on antibiotics (Figure 4b).

**Figure 4.**
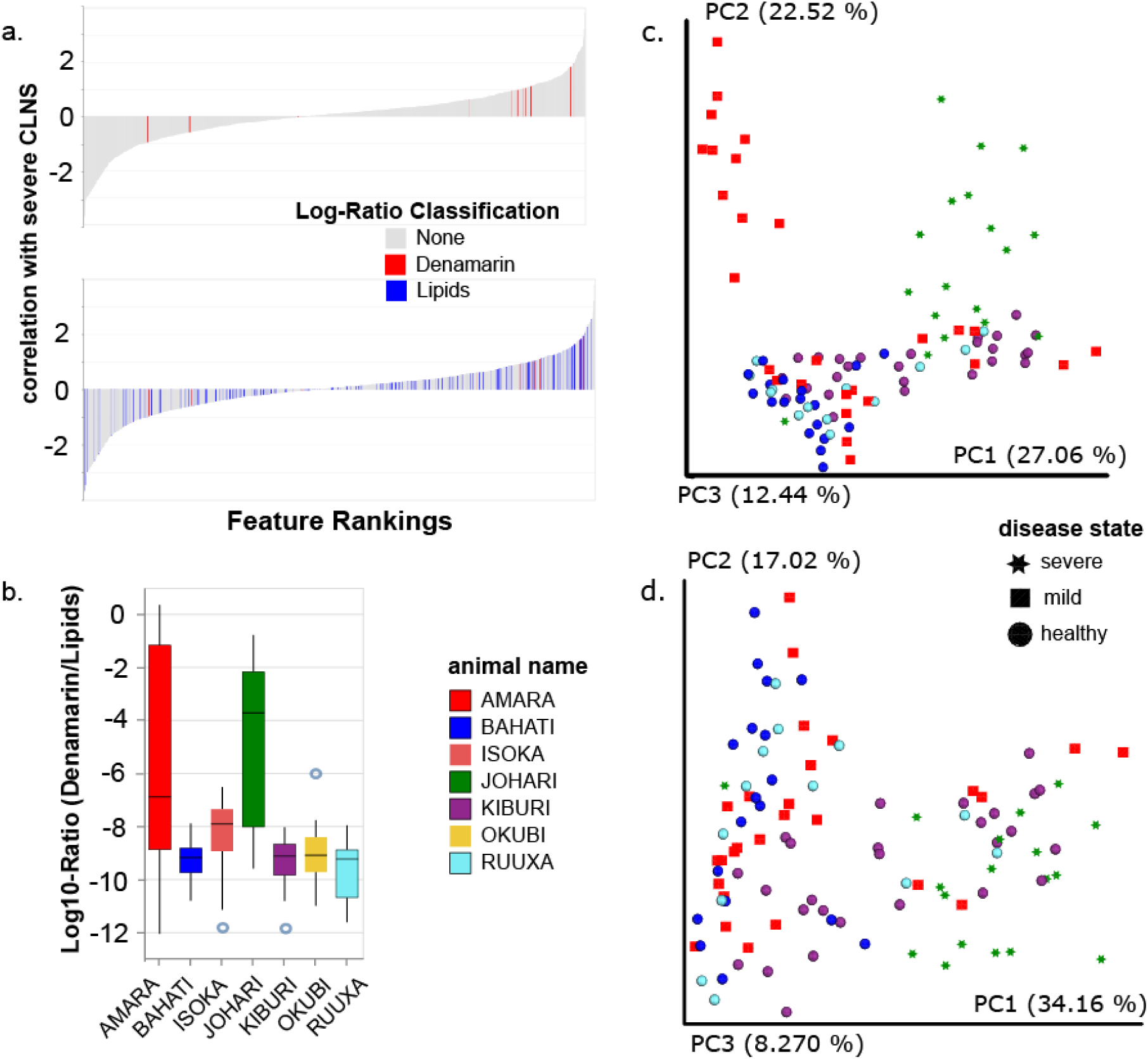
Metabolome-informed filtering of metabolite feature table to remove medication metabolites. Differential abundance analysis of MS features (association with severe liver disease compared to healthy as the reference) visualized with Qurro. Positive values in the rank plots correspond to a positive association with liver disease. **a.** Top: denamarin features highlighted in red; Bottom: Denamarin features in red and lipids as a reference in blue. **b.** Log ratio of denamarin features by the general category lipids plotted by animal. Amara and Johari, both diseased animals, have increased levels of denamarin when compared to the other animals. A PCoA analysis (weighted Unifrac) of 103 samples from WD (5 animals, all no antibiotic exposure), with all features **(c)** show a distinct separation for Amara (red) and Johari (green), while a PCoA analysis (weighted Unifrac) with the features from the metabolism of the supplement denamarin removed **(d)** shows this difference along PC2 was due to this confounder. (Denamarin is defined as the following metabolites: adenine, 5’-Methylthioadenosine, S- (5’-Adenosyl)-L-methionine cation). In d) Johari separates along PC1 from healthy animals, but with some overlap from Kiburi and Amara. Shape designates the disease state, with regards to cheetah liver necrosis syndrome (CLNS); All figures are colored by individual cheetah.

Metabolome data-informed grouping, which was empirically validated, could be used to stratify individuals, to identify criteria of interest, or to flag samples for exclusion from further analysis. In this case, removal of samples from animals exposed to antibiotics, Isoka and Okubi, and recomputation of the PCoA of metabolite data revealed new trends (Figure 4c). Exclusion of samples from Isoka and Okubi (Figure 4c) resulted in no per individual differences along PC1, however, there was a clear separation of Johari (severe disease) and Amara (mild disease), from the remaining healthy individuals along PC2, representing 22.52% of the variance.

We hypothesized that the strong signature from denamarin metabolites was driving this difference in beta diversity and a further iteration of MS-informed analysis was performed (Figure 1). The untargeted metabolomics feature table was filtered to remove metabolite signatures for S-adenosyl methionine, as well as its breakdown products, from the metabolite feature table (red in Figure 4a). Analysis of the modified set of features resulted in decreased separation of metabolite signatures from the cheetahs Johari and Amara (Figure 4d). We observed that along PC1, Johari, with severe CLNS (data points displayed as green stars), was most disparate, while Amara, with a diagnosed mild case (red squares), was not clearly distinguishable from the healthy individuals, as seen in the metagenomic analysis as well.

Although the sample size is small, as is common with studies in captive wildlife, a comparison of clinical data for Amara and Johari (Table S2) support the groupings observed in the final metabolome-informed analyses (Table 4d). Johari had the most progressed case of CLNS, and had liver enzyme levels of alanine aminotransferase (ALT) = 451 and aspartate aminotransferase (AST) = 414 μmol/L on 1/23/18 (critically elevated above the mean reported healthy values ALT = 47 and AST = 93). Amara, who is not clearly distinguished from healthy individuals had elevated ALT and AST levels of 273 and 108 μmol/L in 2017, at the time of diagnosis by liver biopsy, however, on 3/20/18, just after sampling for this study, her levels were within healthy limits at 61 and 13 μmol/L for ALT and AST, respectively. Peripheral bile acid levels were similarly elevated for Johari as well as Amara in 2017 but within normal limits for Amara in 2018 (Table S2). The metabolome profile and presence of Amara’s samples with other healthy individuals is supported by the lower ALT, AST and bile acid levels measured shortly after the sampling time period, indicating a possible recovery back to a more healthy phenotype within this case study.

## 3. Discussion

There are confounders in clinical datasets that cannot be predicted based on available information. Broad untargeted metabolomics gives insights into specific molecules that can reveal unexpected factors that, if not understood, might lead to misinterpretation of the data. In this study, we observed an individual reported not to have been administered antibiotics during the study period but detected antibiotics in their stool. Taking this information into account to generate an MS-informed metadata category for antibiotic use allowed a more informative analysis of the samples. Similarly, upon observation of denamarin metabolites, their signatures could be removed as a result of treatment for liver disease, rather than attributing them as drivers of the condition.

This case study in captive cheetahs illustrates something unique: if the cohort were made up of humans living in a household together, then one individual would ‘correctly’ have self-reported antibiotic use, while the other would have also ‘correctly’ reported not taking antibiotics (or have been monitored by a clinician as not taking antibiotics), but these reports do not accurately reflect antibiotic exposure. MS-informed analyses driven by empirically detected information can therefore reveal elusive information and go beyond even the best (and most intrusive) clinical study. The use of antibiotics by one human household member has been shown to impact the gut microbiota of other members of the same household (10). Although the direct mechanisms of microbiome alteration were not clear in this study, it is reasonable to hypothesize that antibiotic exposure through physical contact with one another, or via household surfaces, may contribute to the carryover of the medications and ultimately impact the gut microbiome. This further raises the question of privacy concerns and whether information from spouses or household members (perhaps even including pets) would also need to be collected for correct interpretation of clinical trials, or if categories such as ‘has anyone in your household taken antibiotics within the last week?’ would be appropriate to include across all studies. The carryover of substances, such as personal care products, medication, and food-derived molecules, from an individual to household surfaces has been well documented (16, 17), however, the ability to transfer antibiotics or indeed other medications from one person to another still remains to be verified.

Identification of confounders that cannot be taken into account initially is a reality of clinical data for captive animals, as we have shown, and is likely also the case for human studies. In the current study, to address the signatures of liver disease, the observation of unexpected antibiotics across numerous samples was very valuable; otherwise Okubi’s samples (no Rx) that contained antibiotics would continue to be included as ‘healthy’, which is true with respect to liver disease, but there is a substantial and clear shift in this cheetah’s microbiome due to antibiotic exposure. As antibiotics are a stronger driver of difference than the disease state, it was appropriate in this case to remove the samples from both animals from the analysis, as they would otherwise be classified as ‘healthy’ because neither suffers from CLNS.

Metabolomics generates a verifiable readout that allows us to reinterpret the study results in the context of the data or information that were actually collected and can be done as an iterative process. Regardless of the origin of such observations, whether it is metadata omission, misreporting, errors in sample collection, or other causes, the fact remains that something is detected where it was not expected. The power of using metabolomic data to reinterpret metadata categorization, impacting all facets of gut microbiome data interpretation, has a critical place in microbiome analyses going forward and virtually any study with samples amenable to MS data collection could benefit from this approach.

## 4. Materials and Methods

### Study design

Between 2015 and 2018, multiple cheetahs from the San Diego Zoo Safari Park collection were diagnosed with Cheetah Liver Necrosis Syndrome (CLNS), via liver biopsy or post mortem exam. A working group including clinical veterinarians, pathologists, nutritionists, and husbandry staff met regularly to develop plans for earlier detection, treatment, disease characterization, and to elucidate the cause of the condition. Fecal samples were obtained from both sick and healthy cheetahs at the Wildlife Discoveries (WD) park. The project was exempt from IACUC review as samples were collected non-invasively as part of a planned, potentially corrective, diet change. Fecal samples were shared between San Diego Zoo Safari Park and UCSD researchers following completion of an External Biomaterials Request in order to explore whether metabolomics and metagenomic data offered a more detailed biochemical picture of ingested materials, both dietary and environmental, beyond the limited nutritional analyses already available.

### Sample handling

Fecal samples were collected each morning daily by trainers at WD for 4 weeks from mid-January 2018 – mid-February 2018. Fecal material was collected on location by animal care staff or transported to the clinical lab at Harter Veterinary Medical Center (HVMC) and sampled by a senior nutrition research associate. Samples were not subjected to extremes of heat or cold (i.e., direct sunlight, vehicle cab, or refrigerator/freezer) during transport. A fecal score was assigned to each individual’s sample. A score of 1 = liquid/watery (runny), 2 = soft (loose, overly moist), 3 = formed (maintains shape), 4 = hard/dry (firm, lacking moisture). Each sample was assigned a unique barcode ID and a label was affixed to the swab tube and a small printed label to the 2ml tube, making sure both samples had the corresponding number for the same sample and animal.

Two separate samples were collected per individual cheetah, a swab of the fecal surface and a small amount of whole feces in a 2mL microcentrifuge tube. The interior of the feces were swabbed with a sterile barcoded cotton swab (BD SWUBE™ Applicator) when possible, or the exterior for smaller samples for sequence analysis. An aliquot of bulk stool was transferred to an empty 2 mL round bottom microcentrifuge tube for mass spectrometric analysis (Qiagen, Hilden, Germany). Samples were frozen at −80 °C within 24 hr of deposition at the San Diego Zoo Institute for Conservation Research (ICR). All samples were transported on dry ice and stored at −80 °C until analysis. All samples were transported on dry ice and subsequently stored at −80 °C until analysis.

All information regarding fecal samples for each animal were recorded into an Excel spreadsheet (Microsoft, Seattle, WA). Information included sample ID, location, date, animal name, institution ID, time feces deposited (if known), time feces collected, time feces sampled, fecal score, any medical / chronic problems (i.e., fecal quality, medications), day and progression of diet transition.

### Metabolomics sample extraction

All fecal samples were dried, weighed and extracted to the same final concentration. Frozen cheetah fecal samples were placed on ice. The lid and rim of each sample tube was cleaned with a Kimwipes™(Kimberly-Clark) moistened with 70% EtOH, to remove spurious fecal material. Samples were dried down overnight using a Labconco CentriVac and either placed at −80 °C for storage, or immediately processed as follows. A clean spatula and tweezers were used to transfer 50-100 mg of dried stool into a new, labeled tube. Exact weights were recorded and a 10-fold volume of cold 50% MeOH in water was added to each tube (ex: 50 mg (stool) + 500 µL (50% MeOH)) (LC-MS grade solvents, Fisher Chemical). The samples were homogenized for 5 minutes at 25 Hz on a tissue homogenizer (QIAGEN TissueLyzer II, Hilden, Germany), and subsequently placed at −20 °C for 15 minutes for methanol extraction. The samples were then centrifuged at max speed (14000 x g) for 15 minutes (Eppendorf Centrifuge 5418; USA). Without disrupting the pellet, 300 µL of the supernatant was transferred into a 96-deep well plate. Samples were sealed and stored at −80 °C.

### Metabolomics data acquisition and processing

Data were collected using a modification of the data dependent acquisition method outlined in (18). Briefly, extracts were dried down, resuspended in 50% MeOH:50% Water (Optima LC-MS grade; Fisher Scientific, Fair Lawn, NJ, USA). Untargeted metabolomics was carried out using an ultra-high-performance liquid chromatography system (UltiMate 3000, Thermo Scientific, Waltham, MA) coupled to a Maxis Q-TOF (Bruker Daltonics, Bremen, Germany) mass spectrometer with a Kinetex C18 column (Phenomenex Torrance, CA, USA). A linear gradient was applied: 0-0.5 min isocratic at 5% B, 0.5-8.5 min 100% B, 8.5-11 min isocratic at 100% B, 11-11.5 min 5% B, 11.5-12 min 5% B; where mobile phase A is water with 0.1% formic acid (v/v) and phase B is acetonitrile 0.1% formic acid (v/v) (LC-MS grade solvents, Fisher Chemical). Electrospray ionization in positive mode was used.

#### MS1 Feature Finding and Data Processing

qToF files (.d) were exported using DataAnalysis (Bruker) as .mzXML files after lock mass correction using hexakis (1H, 1H, 2H-difluoroethoxy) phosphazene (Synquest Laboratories, Alachua, FL), with *m/z* 622.029509. Data quality was assessed by qualitatively evaluating the *m/z* error and retention time of the LC-MS standard solution (i.e. mixture of 6 compounds), which was analyzed at least once in every 96-well plate.

MS1 feature finding was performed on the .mzXML files in MZmine2 (version MZmine-2.37.corr16.4) (Pluskal et al., 2010). The code for this version of mzMINE has been archived with the raw data files available on MassIVE under MSV000082969. The mzMINE parameters used for feature finding are as follows: mass detection (centroid; ms1: 1.5E3; MS2: 90); ADAP Chromatogram builder (min group size in # of scans: 4; group intensity threshold: 5E3; min highest intensity: 2E3; m/z tolerance: 0.001 *m/z* to 20 ppm); chromatogram deconvolution (LMS: chromatographic threshold of 96%, search minimum in RT range (min) of 0.03, minimum relative height of 5%, minimum absolute height of 2E3, min ratio of peak top/edge of 1 and peak duration range (min) of 0 - 2; *m/z* center calculation set to auto; *m/z* range for MS2 scan pairing (Da) of 0.02 and RT range for MS2 scan pairing (min) of 0.15); isotope peaks grouper (*m/z* tolerance set to 0.0015 m/z or 10 ppm; retention time tolerance of 0.05, maximum charge of 3 and representative isotope set to most intense); order peak lists; join aligner (*m/z* tolerance set at 0.0015 *m/z* or 15 ppm; weight for *m/z* of 2; retention time tolerance of 0.2 min; weight for RT of 1. A filter was used such that only features present in at least 2 samples were included. The output was a data matrix of variables (i.e. MS1 features that triggered MS2 scans) by samples, exported for GNPS (.mgf and .csv quant table) and for SIRIUS (.mgf). The MS1 data matrix (MS1 features from quant table) was processed by concatenating the *m/z* and retention time columns from the original MZmine output.

Feature based molecular networking was performed and library IDs were generated using GNPS (14).

The quant table, SIRIUS export, and library identifications from feature based molecular networking were used as inputs for the Qiime2 plugin Qemistree (https://github.com/biocore/q2-qemistree), to perform hierarchical ordering of the untargeted mass spectrometry data. The resultant Qemistree-based feature table can be linked to the original feature number from feature finding (which also links to the GNPS Library IDs) and presents fingerprints that act to merge spectra assigned the same identity. Qemistree also generates a fingerprint-based tree, allowing for tree-based approaches such as UniFrac (19, 20).

#### Applicability of weighted UniFrac (19) for Qemistree abundance analysis

Mass spectrometry feature data represent relative abundances of compounds, not exact counts. Due to this difference, the use of unweighted methods, which only look for presence/absence, are not suitable. Unweighted UniFrac (20) would exaggerate differences between samples if even a small number of new molecules appear in one batch and not the other as frequently occurs due to instrument performance changes during sample processing. Parameters such as gap filling and feature finding parameter settings can further compound these issues. Therefore, weighted UniFrac is used for all comparisons between mass spectral data which were processed using the QIIME 2 plugin q2-qemistree.

QIIME 2 (21) was used within a jupyter notebook environment for Principal Coordinates Analysis. Differential abundance analysis was performed using Calour (or replace with Songbird) (22).

### Sample preparation and sequencing data generation

Shallow shotgun sequencing was performed as previously described (23). In brief, DNA extraction was performed using the Qiagen PowerSoil DNA extraction kit following the Earth Microbiome Project (EMP) standard protocol (24). The Qubit™ dsDNA HS Assay (ThermoFisher Scientific) was used to determine concentration and libraries were prepared from 1 ng of input DNA in a miniaturized Kapa HyperPlus protocol. Libraries were quantified using the Kapa Illumina Library Quantification Kit, pooled and size selected (300-800 bp) using the Sage Science PippinHT. The pooled library was sequenced as a paired-end 150-cycle run on an Illumina HiSeq 4000 mode at the UCSD IGM Genomics Center.Demultiplexed sequences were trimmed and quality filtered using Atropos v 1.1.5, a fork of Cutadapt (25).

#### Sequence data processing

Significant host contamination was expected from the horse, beef, and rabbit-based diet of the cheetahs, as well as host DNA present in the samples. Therefore we identified reads in the quality-filtered reads using Bowtie 2 v2.3.0 (26) with the ‘very-sensitive’ parameter setting against the genomes of *Acinonyx jubatus* (cheetah, isolate: AJU 981 Chewbacca, GCF_001443585.1), *Bos taurus* (cattle, Breed: Hereford, GCF_000003055.6), *Equus caballus* (horse, Breed: thoroughbred, isolate: Twilight, GCF_002863925.1), and *Oryctolagus cuniculus* (rabbit, Breed: Thorbecke inbred, GCF_000003625.3) sequentially.

#### Taxonomy generation and statistical analyses

Host-filtered reads were mapped to the 10,575 genomes selected for phylogenetic reconstruction in the Web of Life project (https://biocore.github.io/wol/data/genomes/) using Bowtie 2 within the alignment pipeline SHOGUN (12) using standard parameters. Bowtie 2 mappings were normalized to distribute reads to individual genomes and the resulting output matrix was filtered to remove reads present at less than 0.01% relative abundance per sample. This filtered matrix was used for weighted UniFrac (19) and unweighted UniFrac (20) beta diversity analysis as well as taxonomic summarization using the Web of Life tree (https://biocore.github.io/wol/data/trees) with QIIME 2 (21). Visualizations were prepared using Emperor (27), and matplotlib (28). PERMANOVA (29) analysis was performed in QIIME 2 v. 2018.11 on metabolite and metagenomic distance matrices. The F-statistic was reported as a measure of effect size. Jupyter notebooks of the analyses are available at http://github.com/knightlab-analyses.

Differential abundance of microbes and metabolites, individually, with regard to different metadata variables were calculated using Songbird (30).

The following formula was used to learn the differentials for the microbes

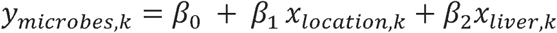

Here, *y*_*microbes*_is the vector of microbe aboundances for a given sample k, *x*_*location,k*_is the location from which the sample was collected from. *β*_1_ is a vector corresponding to the microbe differentials with respect to the sampling location. This represents the log fold differential for each microbe between the sampling locations. *x*_*liver,k*_is an ordinal variable with four possible values CLNS; severe, CLNS, CLNS; mild, healthy. *β*_2_is a vector corresponding to the microbe differentials with respect to the liver health. *β*_0_represents the intercept of the multinomial regression.

The same regression formula was used to learn the differentials for the metabolites.

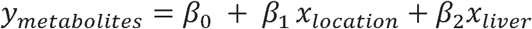

Differentials were visualized in rank plots using Qurro https://github.com/biocore/qurro, while microbe-metabolite co-occurrence probabilities were computed using mmvec https://github.com/biocore/mmvec. These co-occurrence probabilities represent the probability of observing a metabolite given the microbe is observed. These conditional probabilities are estimated through a low-rank approximation with 3 principal axes.

## Data and Code Availability

All data in this study is publically available. Raw and processed shotgun sequencing data is available in Qiita (31) study 11872 (https://qiita.ucsd.edu/study/description/11872) and GNPS (14) using MassIVE (https://massive.ucsd.edu/) ID MSV000082969. The GNPS Networking job is available at https://gnps.ucsd.edu/ProteoSAFe/status.jsp?task=093798dffe2448239410c3d465ef9fea

## Acknowledgements

We thank the keepers and trainers at the San Diego Zoo Safari Park who assisted in sample collection: Jane Kennedy, Dave Gribas, Paula Augusta, Tony Long, Tina Hunter Burnam, Jake Shepherd, Marty Sawin and Janet Rose-Hinostroza, Annette Russell, Shannon Smith-Moritz, Kim Hanley, Larissa Brecht, Ashley Gordon, Kristyn Sargent, Jessica Holland, Kelly Devecchio, Alanna Cappelli. We also thank Katie Kerr and all the staff involved with the care of the animals. Many thanks to Jennifer Damato-Anderson, Jessica Kuang, and Celeste Allaband for assistance pulling clinical records. Many thanks to Anjali Herekar, Caitriona Brennan, Greg Humphrey, Julia Toronczak, Karenina Sanders, MacKenzie Bryant, and Rodolfo Salido-Benitez for shotgun sample preparation. We thank Candace Williams for the introduction to the San Diego Zoo and for helpful comments.

## Author Contributions

JMG, AF conceived the study design.

MG collected samples and coordinated sample collection efforts, contributed to study design.

PD, RK, AS advised data acquisition and analysis.

JMG, CC, KW, SH, RS, AP, MG generated data and metadata. JMG, AT, SH, AS, JM processed and analyzed data.

JMG wrote the manuscript and all authors have approved the final version.

## Supplemental Material

**Table S1**. Top 20 species ranked based on co-occurrence probability with the metabolite soyasaponin I.

**Table S2.** Additional clinical measurements for diseased cheetahs Johari and Amara. **Figure S1.** PcoA plots of metabolome and metagenome showing antibiotic detected metadata characterization.

**Figure S2.** PcoA of shotgun sequence data using unweighted unifrac, with detected categorization.

**Figure S3.** PcoA for shotgun sequence data excluding cheetahs with detectable antibiotics.

**Figure S4.** Taxa bar plots from shotgun sequence data.

