## Supplemental Information for "Metabolome-informed microbiome analysis refines metadata classifications and reveals unexpected medication transfer in captive cheetahs"

Supplementary Materials

**Table S1**. Top 20 species ranked based on co-occurrence probability with the metabolite soyasaponin I. Genome ID assignments were obtained from the Web of Life (https://biocore.github.io/wol/).

| rank | soyasapogenol C (209) | genome ID | NCBI designation |
|---|---|---|---|
| 1 | 5.105 | G000233915 | k__Bacteria; p__Proteobacteria; c__Gammaproteobacteria; o__Xanthomonadales; f__Xanthomonadaceae; g__Pseudoxanthomonas; s__Pseudoxanthomonas spadix |
| 2 | 3.937 | G001375495 | k__Bacteria; p__Actinobacteria; c__Actinobacteria; o__Actinomycetales; f__Actinomycetaceae; g__Flaviflexus; s__Flaviflexus massiliensis |
| 3 | 3.751 | G000227765 | k__Bacteria; p__Proteobacteria; c__Betaproteobacteria; o__Neisseriales; f__Neisseriaceae; g__Neisseria; s__Neisseria wadsworthii |
| 4 | 3.749 | G900113675 | k__Bacteria; p__Firmicutes; c__Bacilli; o__Lactobacillales; f__Carnobacteriaceae; g__Pisciglobus; s__Pisciglobus halotolerans |
| 5 | 3.736 | G001975685 | k__Bacteria; p__Firmicutes; c__Bacilli; o__Lactobacillales; f__Carnobacteriaceae; g__Jeotgalibaca; s__Jeotgalibaca sp. PTS2502 |
| 6 | 3.499 | G000513715 | k__Bacteria; p__Proteobacteria; c__Gammaproteobacteria; o__Vibrionales; f__Vibrionaceae; g__Salinivibrio; s__Salinivibrio socompensis |
| 7 | 3.451 | G000407245 | k__Bacteria; p__Firmicutes; c__Bacilli; o__Lactobacillales; f__Enterococcaceae; g__Enterococcus; s__Enterococcus avium |
| 8 | 3.436 | G001431645 | k__Bacteria; p__Proteobacteria; c__Gammaproteobacteria; o__Xanthomonadales; f__Xanthomonadaceae; g__Stenotrophomonas; s__Stenotrophomonas panacihumi |
| 9 | 3.347 | G000307265 | k__Bacteria; p__Firmicutes; c__Clostridia; o__Clostridiales; f__Oscillospiraceae; g__Oscillibacter; s__Oscillibacter ruminantium |
| 10 | 3.343 | G000418515 | k__Bacteria; p__Firmicutes; c__Bacilli; o__Lactobacillales; f__Lactobacillaceae; g__Lactobacillus; s__Lactobacillus casei |
| 11 | 3.326 | G001425785 | k__Bacteria; p__Proteobacteria; c__Betaproteobacteria; o__Burkholderiales; f__; g__Rhizobacter; s__Rhizobacter sp. Root1221 |
| 12 | 3.235 | G000702265 | k__Bacteria; p__Firmicutes; c__Clostridia; o__Clostridiales; f__Lachnospiraceae; g__Butyrivibrio; s__Butyrivibrio sp. NC2002 |
| 13 | 3.227 | G000174755 | k__Bacteria; p__Bacteroidetes; c__Flavobacteriia; o__Flavobacteriales; f__Flavobacteriaceae; g__Capnocytophaga; s__Capnocytophaga gingivalis |
| 14 | 3.119 | G000015505 | k__Bacteria; p__Proteobacteria; c__Betaproteobacteria; o__Burkholderiales; f__Comamonadaceae; g__Polaromonas; s__Polaromonas naphthalenivorans |
| 15 | 3.029 | G000193225 | k__Bacteria; p__Proteobacteria; c__Betaproteobacteria; o__Burkholderiales; f__Comamonadaceae; g__Verminephrobacter; s__Verminephrobacter aporrectodeae |
| 16 | 2.828 | G000425865 | k__Bacteria; p__Firmicutes; c__Bacilli; o__Lactobacillales; f__Carnobacteriaceae; g__Lacticigenium; s__Lacticigenium naphtae |
| 17 | 2.776 | G000308255 | k__Bacteria; p__Firmicutes; c__Tissierellia; o__Tissierellales; f__Peptoniphilaceae; g__Anaerococcus; s__Anaerococcus pacaensis |
| 18 | 2.735 | G001656215 | k__Bacteria; p__Proteobacteria; c__Gammaproteobacteria; o__Pseudomonadales; f__Moraxellaceae; g__Moraxella; s__Moraxella catarrhalis |
| 19 | 2.729 | G000980895 | k__Bacteria; p__Proteobacteria; c__Alphaproteobacteria; o__Sphingomonadales; f__Sphingomonadaceae; g__Sphingomonas; s__Sphingomonas sp. Ag1 |
| 20 | 2.728 | G000420745 | k__Bacteria; p__Proteobacteria; c__Alphaproteobacteria; o__Rhodobacterales; f__Rhodobacteraceae; g__Pseudorhodobacter; s__Pseudorhodobacter ferrugineus |

**Table S2.** Additional clinical measurements for diseased cheetahs Johari and Amara. Samples taken on 1/23/18, highlighted in green, were during the sampling period. Those samples from Amara highlighted in yellow are referenced in the text. Expected Results (Based on Best Available Match): Type: Min-Max | Mean [Median] N (Animals)

ALT: Global subsp RI: 30-185 | 93 [85] N=361 (113 animals)

AST: Global subsp RI: 22-91 | 47 [44] N=334 (99 animals)

The reference range for total bile acids is 0-6.9umol/L. Moderate to severe elevation (Greater than 30) is generally consistent with hepatic dysfunction.

| Johari | | | |
|---|---|---|---|
| date | ALT | AST | bile acids (μmol/L) |
| 1/16/18 | 497 | 467 | 101.2 |
| 1/23/18 | 451 | 414 | 25.8 |
| 2/27/18 | 414 | 408 | 88.1 |
| 4/5/2018 | 395 | 344 | na |
| 5/1/2018 | 368 | 480 | 30.8 |
| 6/8/18 | 757 | 344 | 39.5 |
| **Amara** | | | |
| date | ALT | AST | bile acids (μmol/L) |
| 1/27/17 | 273 | 108 | 41.1 |
| 12/20/17 | na | na | 1.9 |
| 3/20/18 | 61 | 13 | <0.1 |
| 8/29/18 | 57 | 23 | <1.0 |

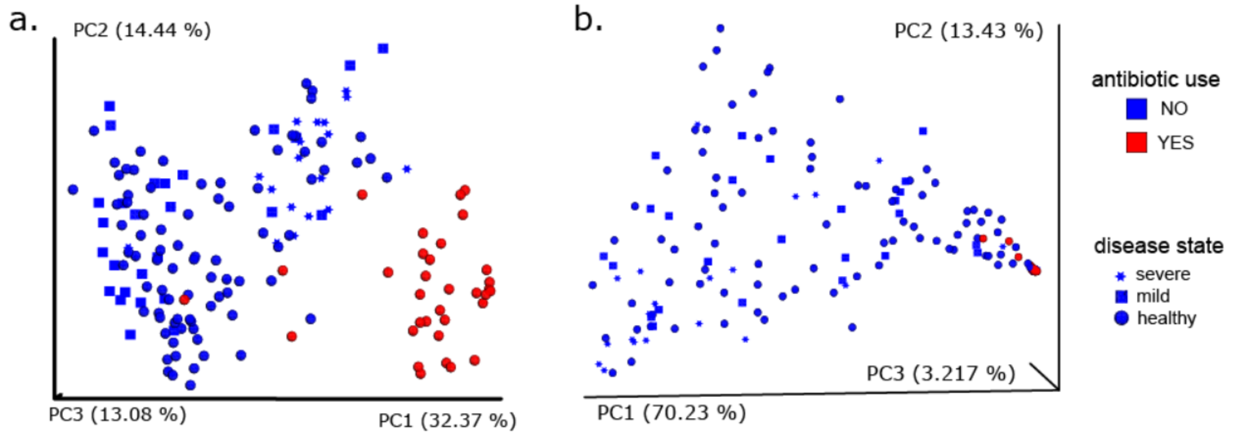

**Figure S1.** Antibiotics are clearly differentiated in the metabolome (a) and metagenome (b) when using MS-informed metadata grouping. Principal coordinate analysis (weighted Unifrac) for metabolome (left) and metagenome (right) for fecal samples from 7 cheetahs housed at Wildlife Discoveries. Data points are colored by MS-informed detected antibiotic use based on empirical evidence (blue: no; red: yes). Shape designates the disease state, with regards to cheetah liver necrosis syndrome (CLNS).

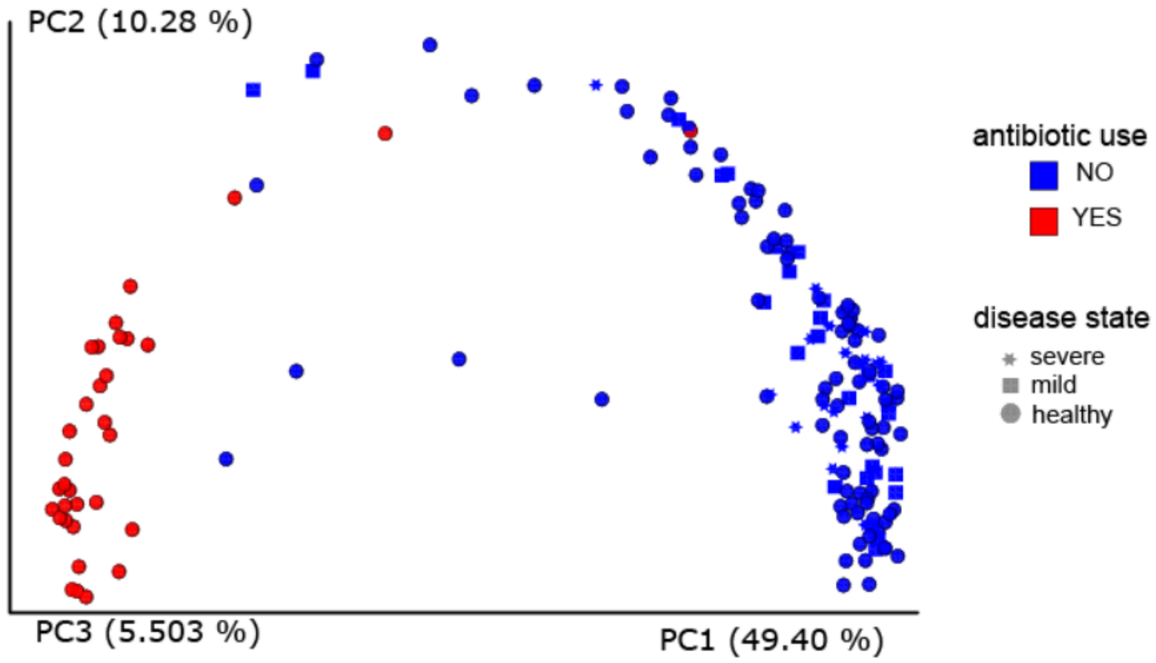

**Figure S2.** Antibiotics are also clearly differentiated in the metagenome when using MS-informed metadata grouping and unweighted unifrac. Data points are colored by MS-informed detected antibiotic use based on empirical evidence (blue: no; red: yes). Shape designates the disease state, with regards to cheetah liver necrosis syndrome (CLNS).

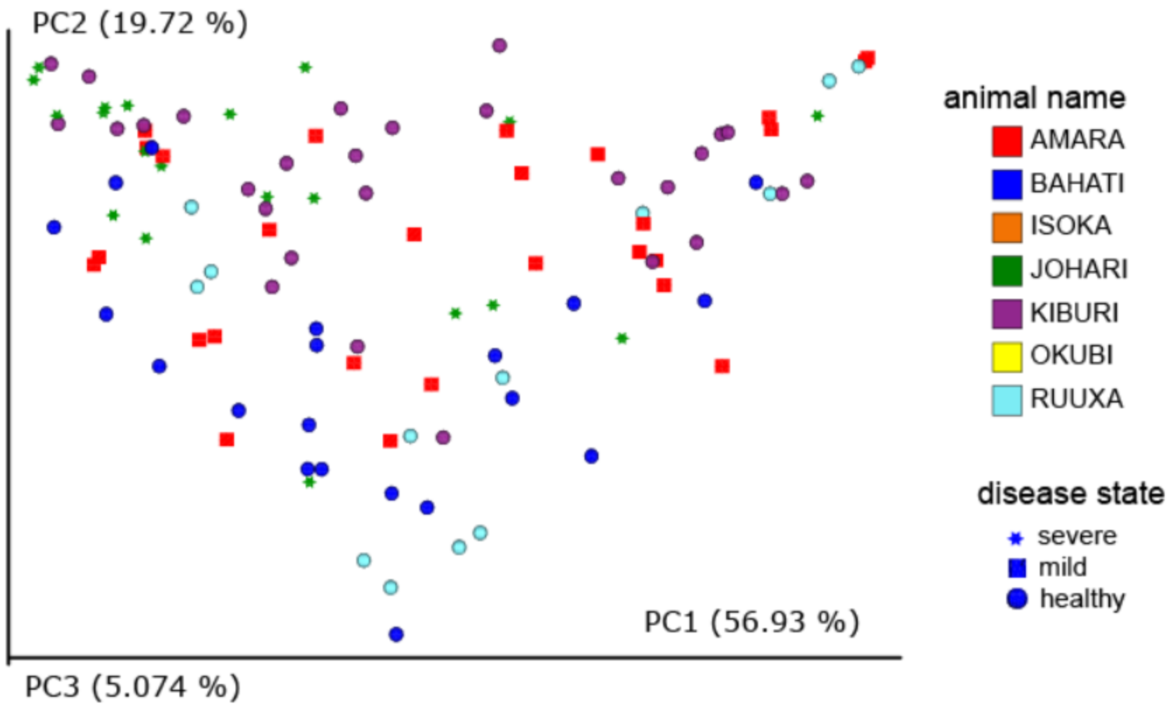

Figure S3. Principal coordinate analysis (weighted UniFrac) for shotgun sequence data for WD animals, excluding Isoka and Okubi, colored by individual. Shape designates the disease state, with regards to cheetah liver necrosis syndrome (CLNS).

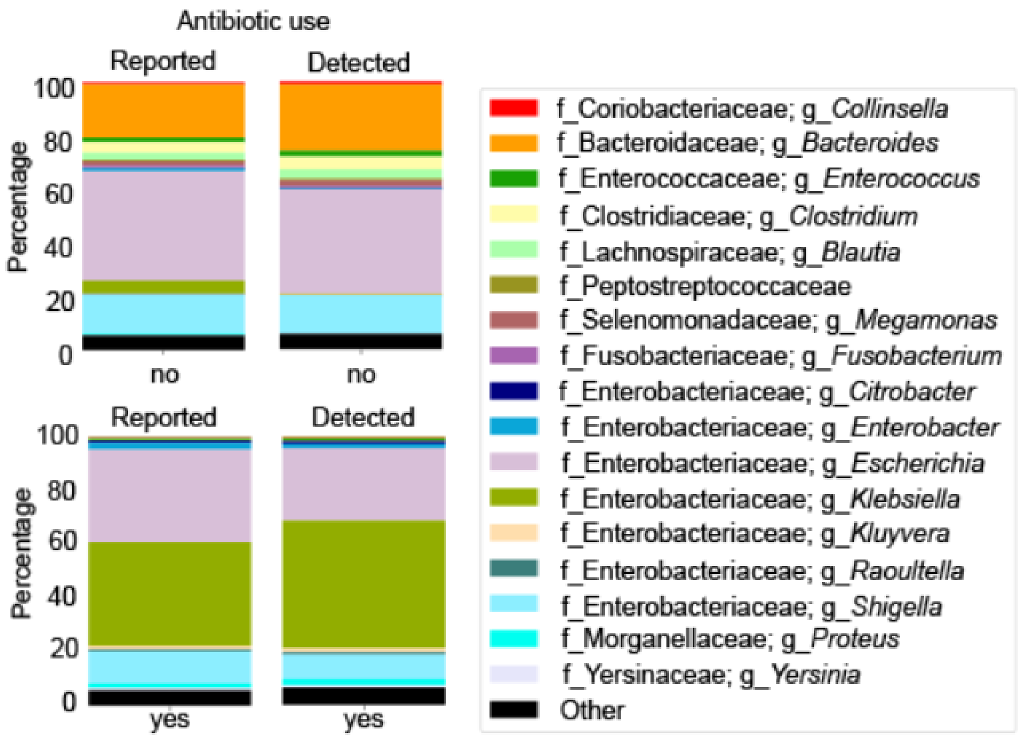

**Figure S4.** Taxa bar plots from shotgun sequence data - showing differences in antibiotic partitioning w/ and w/o metabolome data; left: top 10 genera observed in samples grouped based on original reported metadata; right: top 10 genera observed in samples grouped based on mass spectral analysis (not detected). Klebsiella abundance in the samples with no antibiotic use decreases from reported (4.8%) to empirically detected (0.27%). Other is a summation of the remaining taxa.
